# Allelic chromatin structure primes imprinted expression of *Kcnk9* during neurogenesis

**DOI:** 10.1101/2023.06.09.544389

**Authors:** Daniel Loftus, Bongmin Bae, Courtney M. Whilden, Amanda J. Whipple

## Abstract

Differences in chromatin state inherited from the parental gametes influence the regulation of maternal and paternal alleles in offspring. This phenomenon, known as genomic imprinting, results in genes preferentially transcribed from one parental allele. While local epigenetic factors such as DNA methylation are known to be important for the establishment of imprinted gene expression, less is known about the mechanisms by which differentially methylated regions (DMRs) lead to differences in allelic expression across broad stretches of chromatin. Allele-specific higher-order chromatin structure has been observed at multiple imprinted loci, consistent with the observation of allelic binding of the chromatin-organizing factor CTCF at multiple DMRs. However, whether allelic chromatin structure impacts allelic gene expression is not known for most imprinted loci. Here we characterize the mechanisms underlying brain-specific imprinted expression of the *Peg13-Kcnk9* locus, an imprinted region associated with intellectual disability. We performed region capture Hi-C on mouse brain from reciprocal hybrid crosses and found imprinted higher-order chromatin structure caused by the allelic binding of CTCF to the *Peg13* DMR. Using an *in vitro* neuron differentiation system, we show that on the maternal allele enhancer-promoter contacts formed early in development prime the brain-specific potassium leak channel *Kcnk9* for maternal expression prior to neurogenesis. In contrast, these enhancer-promoter contacts are blocked by CTCF on the paternal allele, preventing paternal *Kcnk9* activation. This work provides a high-resolution map of imprinted chromatin structure and demonstrates that chromatin state established in early development can promote imprinted expression upon differentiation.

## Introduction

Prior to fertilization, the maternal and paternal genomes differ in multiple aspects of chromatin state, reflecting the unique processes of oogenesis and spermatogenesis.^1,2,3^ While most of these differences, which include DNA methylation, histone modifications, and higher-order chromatin structure, are equalized by epigenetic reprogramming in early embryogenesis, some withstand this process and are maintained throughout development.^4,5^ These long-term imbalances in chromatin state between the parental alleles, known as genomic imprinting, result in parent-of-origin specific differences in transcriptional levels at approximately 200 genes in placental mammals.^6,7^ While imprinted expression is found across many developmental timepoints and tissues, it is especially prevalent during early embryogenesis and in the developing and adult brain,^8^ and many neurodevelopmental disorders are associated with the dysregulation of imprinted genes.^9,10^

An early observation regarding genomic imprinting was that imprinted genes are not randomly distributed across the genome, but rather tend to occur in clusters.^11^ These large genomic clusters, often exceeding a megabase in size, are linked to the presence of differentially methylated regions (DMRs) known as imprint control regions (ICRs) that are typically a few kilobases or less.^12^ ICRs are resistant to the global wave of DNA methylation erasure that occurs in the early embryo^13,14^ and provide the fundamental basis for imprinted gene expression in adult tissues. Studies examining how ICRs influence the expression of distal genes, sometimes hundreds of kilobases away, have yielded valuable insights into repressive long non-coding RNAs (lncRNAs)^15,16,17,18^ and allele-specific insulators.^19,20,21^ However, the mechanisms by which ICRs influence the expression of distal genes across most imprinted clusters are not fully understood.

The growing use of advanced technologies such as Hi-C is yielding new appreciation for higher-order chromatin organization and its relationship to gene regulation.^22^ Chromatin folding into topologically associated domains (TADs) mediates high frequency contacts between cis-regulatory elements like enhancers and promoters.^23^ Among the most important determinants of TAD structure in mammalian cells is the DNA-binding protein CTCF, which is frequently found at TAD boundaries and can insulate promoters from surrounding enhancers.^23,24^ While many individual examples of the importance of CTCF-mediated chromatin structure on gene expression exist,^25,26,27^ severe ablation of native chromatin structure through the acute depletion of CTCF produces only modest changes in global gene expression,^28,29^ leading to questions about the precise role of CTCF and higher-order chromatin structure more broadly in regulating transcription.

Studies of genomic imprinting have played an important part in our understanding of CTCF-mediated chromatin folding. Most notably, the *H19-Igf2* locus has long served as a critical model system for CTCF function, higher-order chromatin structure, and enhancer-mediated gene regulation.^30,31,32^ Early studies of the *H19-Igf2* locus found that CTCF binds only the unmethylated maternal ICR and plays multiple roles in regulating imprinted expression at the locus.^33^ First, allelic binding of CTCF leads to allelic higher-order chromatin structure at the *H19- Igf2* locus, thereby regulating imprinted gene expression through parent-of-origin differences in enhancer-promoter contacts.^19,20^ Second, allelic CTCF binding to the maternal allele is essential for maintaining the unmethylated state of the ICR and surrounding secondary DMRs.^34^ Third, CTCF is important for maintaining allelic differences in histone modifications at the locus.^35^ Allelic CTCF binding has since been shown to occur at many imprinted loci, which often correlates with allele-specific chromatin folding patterns. For example, allelic higher-order chromatin structure has been implicated in regulating imprinted expression at the murine *Dlk1-Dio3* and *Grb10-Ddc* loci.^20,21^ Additionally, recent work has revealed that allelic chromatin structure is present at multiple imprinted loci in human.^36^ However, whether this allelic chromatin folding is functionally important for allelic gene expression has not been determined at most imprinted clusters.

Here we characterize the role of allelic CTCF binding and chromatin structure at the *Peg13-Kcnk9* imprinted locus, an important gene cluster with brain-specific imprinted expression.^37,38,39^ Genetic mutations within this imprinted cluster are associated with intellectual disability,^40,41,42^ most notably Birk-Barel syndrome, which is caused by maternal inheritance of a missense mutation in the potassium leak channel *Kcnk9*.^43^ We performed region capture Hi-C on brain tissue from reciprocal hybrid mouse crosses and found differences in TAD structure, insulation, and enhancer-promoter contact frequency between the two parental alleles due to allelic CTCF binding at the *Peg13* DMR. Deletion of the paternal CTCF binding sites in an *in vitro* neuron differentiation system led to a maternalization of paternal chromatin structure and accompanying loss of *Peg13-Kcnk9* locus imprinted expression. Moreover, we found that allelic TAD structure precedes imprinted expression at the locus, and that pre-existing allelic chromatin structure is sufficient to induce maternal-specific expression of *Kcnk9* upon activation of distal enhancers. This work contributes to our understanding of how higher-order chromatin structure can regulate tissue-specific imprinted expression and has broad implications for the role of pre-existing chromatin structure in developmental gene expression.

## Results

### Chromatin structure at the *Peg13*-*Kcnk9* locus is imprinted

The *Peg13-Kcnk9* locus is composed of the lncRNA gene *Peg13* and the protein-coding genes *Kcnk9*, *Trappc9*, *Chrac1*, and *Ago2* (Fig. 1A). Imprinted expression of the locus has been observed in brain tissue but not in body tissues.^8,37,38,39^ More specifically, in mouse brain tissue, *Kcnk9* is expressed exclusively from the maternal allele, *Peg13* is expressed exclusively from the paternal allele, and *Trappc9*, *Chrac1*, and *Ago2* exhibit a maternal bias. Imprinted expression of *Kcnk9* and *Peg13* have also been observed in human brain.^39^ In body tissues, *Kcnk9* and *Peg13* are transcriptionally silent, and *Trappc9*, *Chrac1*, and *Ago2* are expressed in a biallelic manner. Currently, it is not known how imprinted expression of these genes is achieved, or why imprinted expression is exclusively observed in the brain. A maternally methylated DMR overlaps the promoter of *Peg13*^4^ (Fig. 1A) and is the putative ICR controlling imprinted expression of the entire locus.^39,44,45^ Analysis of publicly available CTCF ChIP-seq datasets from mouse brain showed paternal-specific CTCF binding at the *Peg13* DMR (Fig. 1A),^46^ consistent with the known antagonistic relationship between DNA methylation and CTCF binding.^47,48,49^

**Figure 1.**
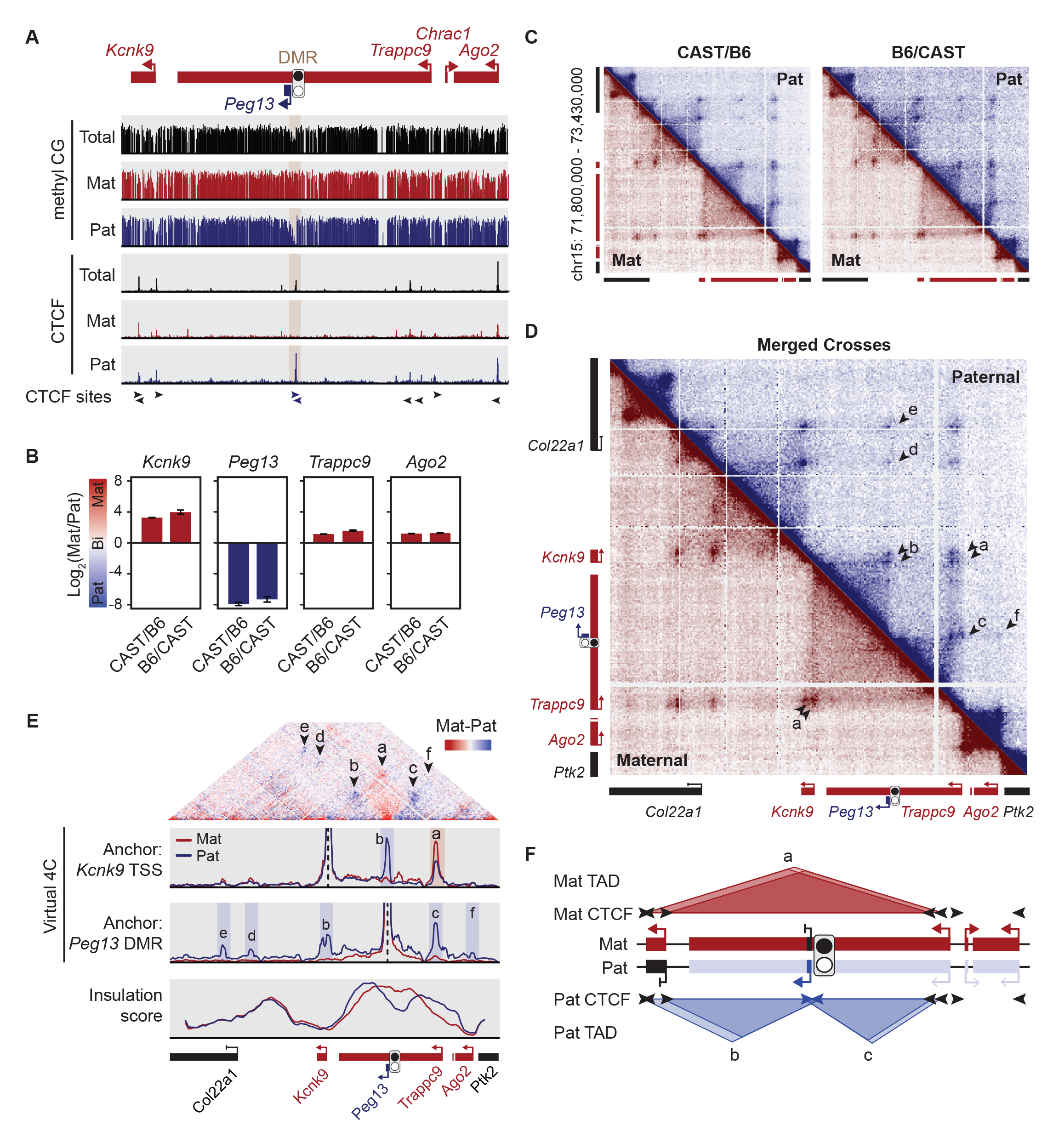
Chromatin structure at the *Peg13-Kcnk9* locus is imprinted. **A.** Allelic CpG methylation (reanalyzed from Xie et al., 2012^4^) and allelic CTCF binding (reanalyzed from Prickett et al., 2013^46^) at the *Peg13-Kcnk9* locus in mouse brain. **B.** Ratio of maternal to paternal expression in brain tissue from reciprocal hybrid mouse crosses as measured by RT-ddPCR. Note that biallelic expression is when y = 0. Mean ± SEM (n=3). **C.** Allelic contact maps from region capture Hi-C of reciprocal hybrid mouse brain. **D.** Merged allelic contact maps combining both crosses from (C). Arrowheads indicate contacts of interest. **E.** (Top) Hi-C subtraction plots (maternal – paternal) showing allelic biased contacts. (Middle) Virtual 4C analysis of the region capture Hi-C datasets anchored at the *Kcnk9* TSS or the *Peg13* DMR. Contacts of interest are highlighted. (Bottom) Allelic insulation score analysis. **F.** Diagram summarizing imprinted chromatin structure and expression at the *Peg13-Kcnk9* locus in brain. See also Figure S1.

In order to determine if allelic CTCF binding at the *Peg13* DMR results in allelic chromatin structure, we generated reciprocal mouse crosses of the distantly related mouse strains *M.m.musculus* (B6) and *M.m.castaneus* (CAST). Leveraging single-nucleotide polymorphisms (SNPs) to distinguish between the B6 and CAST genomes, we first developed an allele-specific droplet-digital PCR (ddPCR) assay (Fig. S1) and observed the expected imprinted expression of *Kcnk9*, *Peg13*, *Trappc9*, and *Ago2* in reciprocal mouse brain tissue (Fig. 1B). *Chrac1* does not contain any exonic SNPs and was therefore excluded from analysis. We then performed region capture Hi-C on brain tissue from reciprocal hybrid crosses using biotinylated capture probes to 1.5 Mb of chromatin at the *Peg13-Kcnk9* imprinted locus and neighboring genes. Hi-C reads containing strain-specific SNPs were used to generate allelic contact maps. Allelic contact maps showed clear differences in the chromatin contacts between the maternal and paternal genomes across the *Peg13*-*Kcnk9* locus (Fig. 1C, D). Similar parent-of-origin effects were observed in both crosses (Fig. 1C), and the contact maps from reciprocal crosses were therefore merged for further analysis (Fig. 1D). On the maternal allele a single TAD crossing the DMR predominates, anchored by a strong contact between the *Kcnk9* transcription start site (TSS) and a distal region in an intron of *Trappc9* (Fig. 1D, arrowhead ‘a’). On the paternal allele, this TAD is less pronounced and two additional smaller TADs are observed that are anchored at the CTCF-bound *Peg13* DMR (Fig. 1D, arrowheads ‘b’ and ‘c’). Long-range, paternal-specific contacts anchored at the DMR are also seen (Fig. 1D, arrowheads ‘d’ and ‘e’), as well as a weak paternal-specific contact between the DMR and the *Ago2* TSS (Fig. 1D, arrowhead ‘f’).

The parental bias in contacts across the imprinted locus can be further visualized as allelic subtraction maps (maternal minus paternal reads) (Fig. 1E, top). This view reveals that contacts crossing the *Peg13* DMR are diminished on the paternal allele, reflecting the insulating effect of paternal-specific CTCF binding. We additionally performed virtual 4C analysis of the allelic Hi-C data from two anchor points, the *Kcnk9* TSS and the DMR (Fig. 1E, middle). Using the *Kcnk9* TSS as the anchor, we confirmed a maternally biased contact with the *Trappc9* intronic region (arrowhead ‘a’) and a paternal-specific contact with the DMR (arrowhead ‘b’). Using the DMR as the anchor, we confirmed that all major contacts made by the DMR are paternal-specific (arrowheads ‘c’-‘f’), thereby demonstrating the strong effect of paternal-specific CTCF binding at the unmethylated DMR. Finally, allelic insulation score analysis highlights the paternal-specific TAD boundary at the *Peg13* DMR (Fig. 1E, bottom). These results provide a high-resolution view of imprinted chromatin structure at the *Peg13-Kcnk9* locus (Fig. 1F) and are consistent with allelic CTCF binding at the *Peg13* DMR as the basis of the structural differences between the two parental alleles.

### Imprinted chromatin structure precedes imprinted expression at the *Peg13*-*Kcnk9* locus

To investigate the mechanisms regulating brain-specific imprinted expression at the *Peg13-Kcnk9* locus, we implemented an *in vitro* neuron differentiation system from hybrid ESCs which is amenable to allelic measurements and rapid genetic perturbations.^50^ Briefly, mouse embryonic stem cells (ESCs) were derived from a B6 x CAST cross. Hybrid ESCs were then differentiated into induced neurons (iNs) by doxycycline-inducible expression of *Ngn2* for 6-12 days. Successful differentiation was noted by down-regulation of the pluripotency genes *Nanog* and *Oct3/4* and upregulation of the neuronal marker *Tubb3* (Fig. 2A).

**Figure 2.**
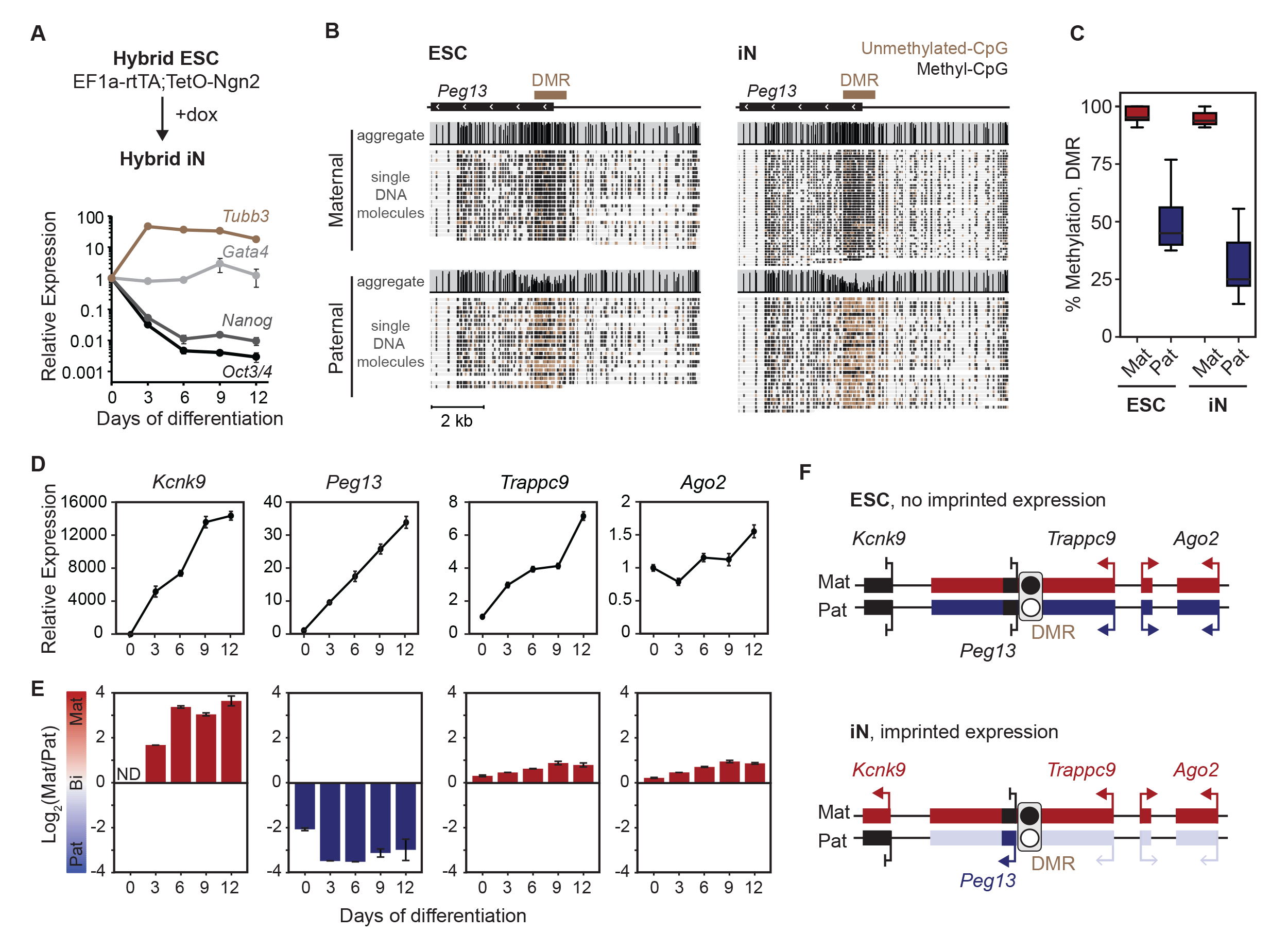
Imprinted expression of the *Peg13*-*Kcnk9* locus is acquired during neuron differentiation. **A.** RT-qPCR of pluripotency and differentiation marker genes during *in vitro* neuron differentiation. **B.** Targeted nanopore sequencing of native genomic DNA to measure allelic DNA methylation levels at the *Peg13* DMR in ESCs and iNs. **C.** Quantification of average methylation levels at individual CpGs at the DMR from (C). **D.** Relative total expression levels of the genes in the *Peg13-Kcnk9* locus during differentiation as measured by RT-qPCR. Mean ± SEM (n=3). **E.** RT-ddPCR during neuron differentiation. Mean ± SEM (n=3). ND, not detected. **F.** Diagram summarizing allelic DMR methylation and gene expression patterns in ESCs and iNs. See also Figure S2.

We first characterized allelic DNA methylation and gene expression patterns at the *Peg13-Kcnk9* locus during neuron differentiation. Targeted nanopore sequencing of native genomic DNA in ESCs and iNs at the *Peg13* DMR showed >90% methylation on the maternal allele and 25-50% methylation on the paternal allele (Fig. 2B,C), a finding which was corroborated using bisulfite Sanger sequencing (Fig. S2A). RT-qPCR showed that *Kcnk9* and *Peg13* are robustly upregulated during neuron differentiation, with modest upregulation of *Trappc9* and *Ago2* (Fig. 2D). ddPCR showed that the expression of *Kcnk9* is strongly maternal in iNs and *Peg13* is strongly paternal (Fig. 2E). *Trappc9* and *Ago2* are bi-allelically expressed in ESCs and acquire a maternal bias upon neuron differentiation (Fig. 2E). The parental bias in expression of all four genes in iNs is similar to mouse brain tissue. These results show that imprinted expression of the *Peg13-Kcnk9* locus is acquired during neuron differentiation in the absence of changes in methylation at the DMR (Fig. 2F). This, therefore, provides a good model system to interrogate the mechanistic basis of cell type-specific imprinted expression.

We next examined how chromatin structure may change during neuron differentiation. First, we performed CTCF ChIP and found that CTCF binds to the *Peg13* DMR in a paternal-specific manner in both ESCs and iNs (Fig. 3A, B). We then performed region capture Hi-C before and after neuron differentiation. We observed imprinted chromatin structure in both ESCs and iNs, and major allelic contacts observed *in vivo* were also seen *in vitro* (Fig. 3C, D). The contact maps are largely similar between ESCs and iNs, with iNs exhibiting some strengthening of existing contacts at nearby CTCF sites (double arrowheads ‘a’ and ‘b’) and gain of additional long-range contacts between the paternal DMR and regions outside the imprinted domain (arrowheads ‘d’ – ‘f’). These results show that imprinted chromatin structure is already established in ESCs and precedes imprinted gene expression at the *Peg13-Kcnk9* locus. This suggests that higher-order chromatin structure may prime the locus for imprinted expression prior to neurogenesis.

**Figure 3.**
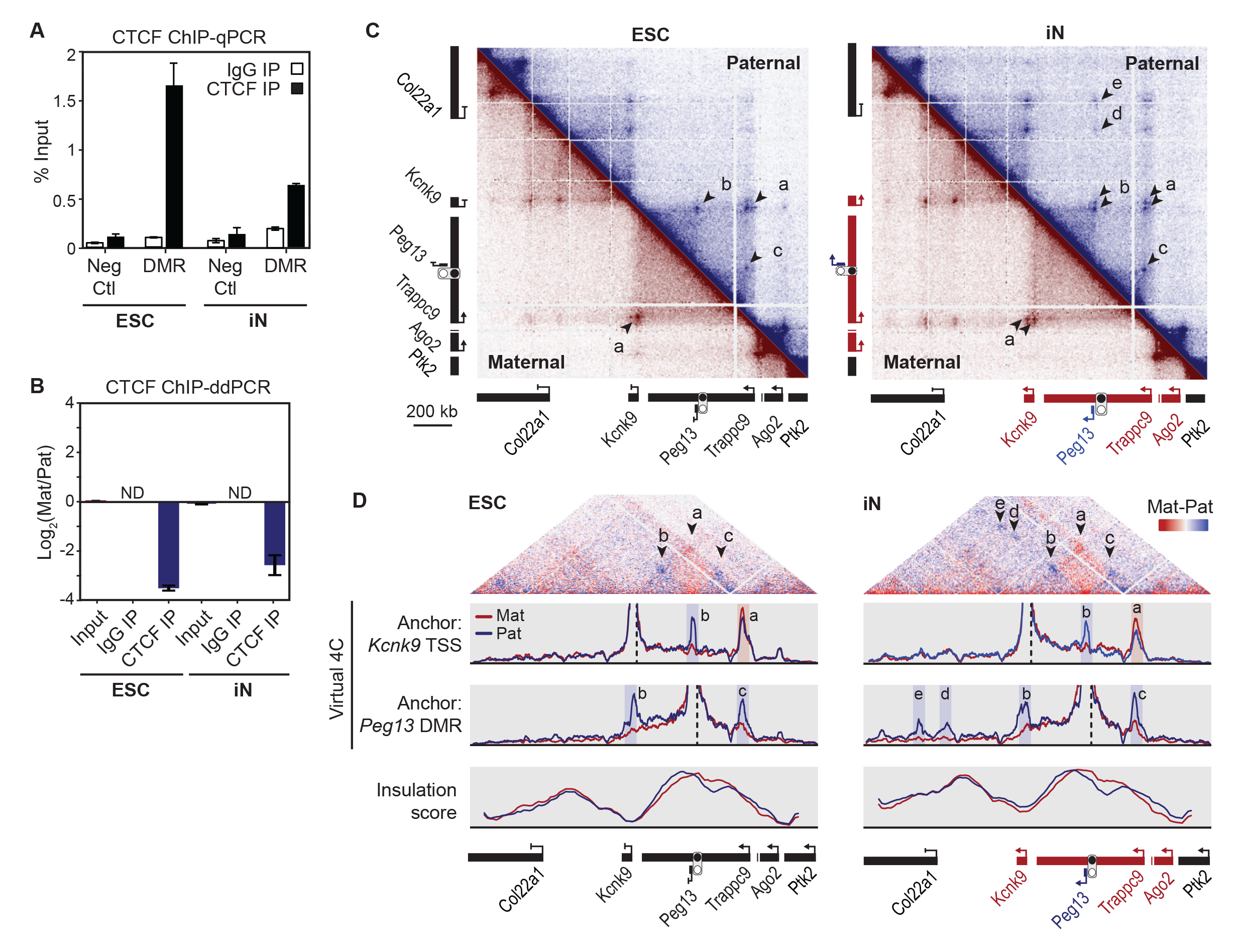
Imprinted chromatin structure precedes imprinted expression at the *Peg13- Kcnk9* imprinted locus. **A.** CTCF ChIP-qPCR at the *Peg13* DMR and a non-CTCF bound control region (Neg Ctl) in ESCs and iNs. Mean ± SEM (n=3). **B.** ChIP-ddPCR for the allelic ratio of CTCF binding in ESCs and iNs. Mean ± SEM (n=3). ND, not detected. **C.** Allelic contact maps from region capture Hi-C in ESCs (left) and iNs (right). Arrowheads indicate contacts of interest. **D.** (Top) Hi-C subtraction plots (maternal – paternal) showing allelic biased contacts. (Middle) Virtual 4C analysis of the region capture Hi-C datasets anchored at the *Kcnk9* TSS or the *Peg13* DMR. Contacts of interest are highlighted. (Bottom) Allelic insulation score analysis.

### The CTCF region of the *Peg13* DMR is essential for imprinted expression

The *Peg13* DMR consists of approximately 2 kb of genomic sequence overlapping the *Peg13* promoter^12^. Given that this is the only DMR that has been identified in the genomic region it is presumed to be the locus ICR, but the mechanism(s) by which the DMR may regulate the allelic expression of the surrounding protein-coding genes is not known. Within the DMR there are two CTCF motifs located approximately 300 bp apart, one in the forward orientation and one in the reverse orientation (Fig. 4A). We hypothesized that the CTCF-bound region of the DMR may be the region critical for imprinted expression.

**Figure 4.**
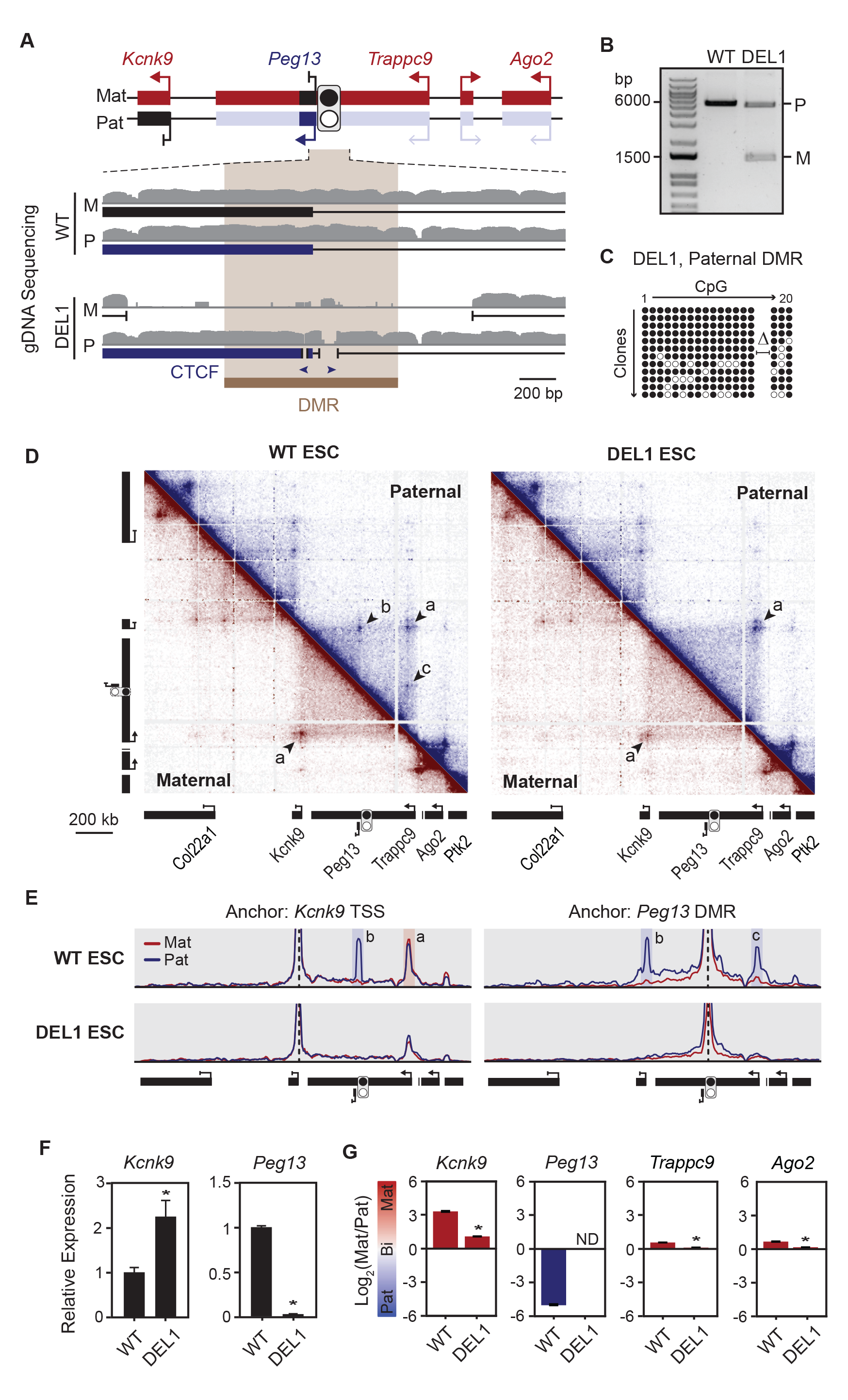
The CTCF region of the *Peg13* DMR is essential for imprinted expression. **A.** Diagram of the *Peg13* DMR depicting the CRISPR-mediated deletions of DEL1. Genomic tracks show single-end mapping of Hi-C genomic DNA. **B.** Agarose gel showing PCR products from the clonal ESC line using primers spanning the *Peg13* DMR. Deletion products were validated by Sanger sequencing to determine parent-of-origin using strain-specific SNPs. **C.** Bisulfite Sanger-sequencing across the *Peg13* DMR in DEL1 ESCs indicating methylated (closed circles) and unmethylated (open circles) CpGs. Two CpGs overlapping the deleted CTCF site are indicated with Δ. **D.** Allelic contact maps from region capture Hi-C of WT and DEL1 ESCs. Arrowheads indicate contacts of interest. **E.** Virtual 4C analysis of region capture Hi-C from WT and DEL1 ESCs. **F.** Total expression measured by RT-qPCR of WT and DEL1 iNs. Mean ± SEM (n=3). **G.** Allelic expression measured by RT-ddPCR of WT and DEL1 iNs. Mean ± SEM (n=3). *p ≤ 0.05, calculated using unpaired two-tailed Student’s t-test. See also Figure S3.

If CTCF binding is necessary for neuron-specific imprinted expression, then deletion of the paternal DMR CTCF sites should result in a maternal-like expression pattern from the paternal allele (i.e., equal gene expression between the two alleles). We used CRISPR-Cas9 and multiplexed sgRNAs to generate two clonal paternal DMR CTCF deletion lines – DEL1 has an 18 bp deletion overlapping the reverse CTCF motif and a 180 bp deletion overlapping the forward CTCF motif (Fig. 4A, B), and DEL2 has a single 348 bp deletion, including all of the forward CTCF site and most of the reverse CTCF site (Fig. S3A, B). Both DEL lines also had large deletions on the maternal allele of ∼1300 bp (DEL1) and ∼4000 bp (DEL2) (Fig. 4A, B; S3A, B).

We asked if deleting the paternal CTCF sites at the *Peg13* DMR had an effect on the structure of the paternal allele. Bisulfite treatment followed by Sanger sequencing of DEL1 revealed that the paternal allele gained methylation upon deletion of the CTCF binding sites (Fig. 4C), suggesting that CTCF maintains the unmethylated state on the paternal allele. Region capture Hi-C on WT and DEL ESCs revealed that loss of paternal CTCF led to a maternalization of chromatin structure on the paternal allele, with an absence of paternal-specific sub-TADs and a strengthening of the long-range contacts crossing the *Peg13* DMR (Fig. 4D, E; S3B, C). Deletion of the methylated maternal ICR had no effect on the chromatin structure of the maternal allele. This demonstrates that the CTCF region of the *Peg13* DMR is critical for imprinted chromatin structure, and is consistent with previous results showing that the human *Peg13* DMR has insulator activity.^39^

We then differentiated WT and DEL ESCs to iNs and measured total and allelic expression at the *Peg13-Kcnk9* locus (Fig. 4F, G). The DEL2 ESCs lost pluripotency and failed to properly differentiate and therefore gene expression analysis was only performed in DEL1 iNs. *Peg13* expression was strongly reduced in DEL1 (Fig. 4F), reflecting the deletion of the *Peg13* TSS. *Kcnk9* expression increased approximately two-fold, consistent with an activation of paternal *Kcnk9* (Fig. 4F). The neuronal allelic biases in *Kcnk9*, *Trappc9*, and *Ago2* were significantly reduced (i.e., trended toward biallelic expression) (Fig. 4G). *Kcnk9* retained a slight maternal bias, which may reflect residual secondary epigenetic effects or additional roles for the non-deleted regions of the DMR. These results demonstrate that the *Peg13* DMR acts as the locus ICR, and that the CTCF-bound region is essential for imprinted chromatin structure and imprinted expression of the *Peg13*-*Kcnk9* cluster as well as for maintaining the unmethylated state of the paternal *Peg13* DMR.

### The *Peg13* lncRNA is not required for imprinted expression

Allelic lncRNAs have been shown to localize to their site of transcription and act *in cis* to repress surrounding genes at other imprinted loci.^15,16,51,52^ We wanted to determine if the *Peg13* lncRNA may mediate repression of neighboring protein-coding genes on the paternal allele. The onset of *Peg13* expression on the paternal allele is correlated with the transition to maternal-biased expression of the surrounding genes during neurogenesis (Fig. 2D, E), and deletion of the *Peg13* TSS in DEL1 cells was correlated with loss of maternal-biased expression of these genes. Additionally, *Peg13* has been shown to interact with the PRC2 complex^53^, which is responsible for deposition of the repressive H3K27me3 histone modification, further supporting this model.

We first determined the subcellular localization of *Peg13* in mouse brain by single molecule fluorescent *in situ* hybridization (smFISH). In contrast to other cis-repressive lncRNAs,^16,54,55^ *Peg13* was cytoplasmic and did not form a large focus near its site of transcription (Fig. 5A). Our observation is consistent with previously published results regarding *Peg13* localization in neural stem cells.^56^ Next, we tested if knockdown of *Peg13* in iNs would result in a loss of imprinted expression of the surrounding protein-coding genes. Treatment of iNs with antisense oligonucleotides (ASOs) against *Peg13* led to a ∼70% knockdown of *Peg13* and a ∼50% knockdown of *Trappc9* (Fig. 5B). The entire *Peg13* sequence is contained in an intron of *Trappc9* and therefore the ASOs are fully complementary to the *Trappc9* pre-mRNA. There was no change in either the total or allelic expression of *Kcnk9* or *Ago2* (Fig. 5B, C). These data suggest that *Peg13* is not required for imprinted expression of *Kcnk9* or *Ago2* in iNs. Furthermore, the loss of imprinting observed in DEL1 iNs is likely not due to deletion of the *Peg13* TSS.

**Figure 5.**
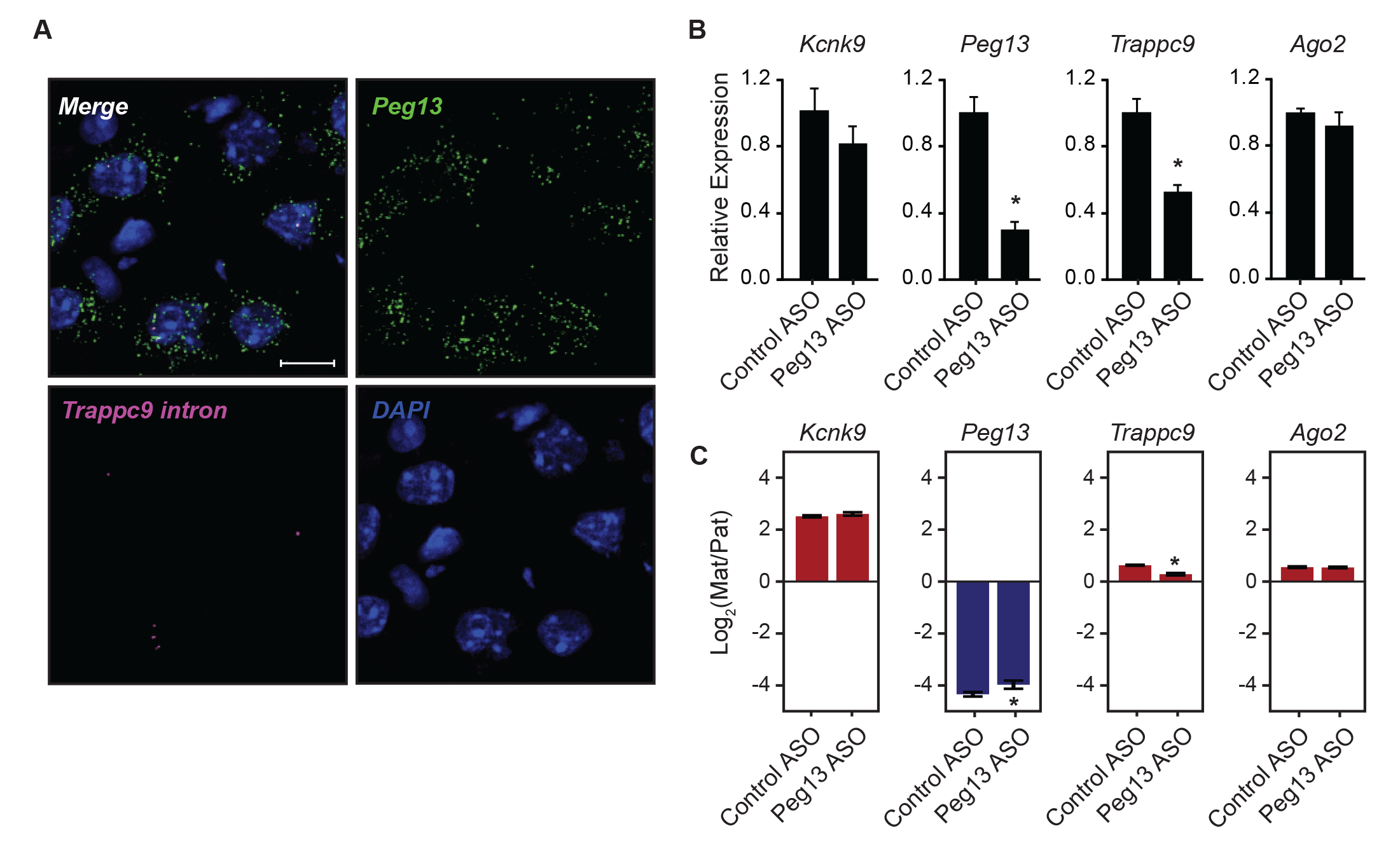
The *Peg13* lncRNA is not required for imprinted expression. **A.** Subcellular localization of *Peg13* in mouse brain sections measured using smFISH probes to *Peg13* and the neighboring intronic region of *Trappc9*. Scale bar = 10 µm. **B.** Total expression measured by RT-qPCR following ASO treatment in iNs. Mean ± SEM (n=3). **C.** Allelic expression measured by RT-ddPCR following ASO treatment in iNs. Mean ± SEM (n=3). *p ≤ 0.05, calculated using unpaired two-tailed Student’s t-test.

### Pre-existing allelic chromatin structure primes imprinted expression of *Kcnk9* upon enhancer activation

We next sought to determine the role of imprinted chromatin structure in the maternal-specific expression of *Kcnk9*. The *Kcnk9* TSS forms maternally-biased contacts that cross the DMR. Potential regulatory elements located downstream of the DMR (with respect to *Kcnk9*) are insulated from *Kcnk9* on the paternal allele by the CTCF-bound DMR, making them strong candidates for regulation of *Kcnk9* imprinted expression. Analysis of publicly available ChIP-seq datasets from mouse tissues suggests that there are at least two tissue-specific, maternal enhancers in this region, which we called E1 and E2 (Fig. 6A). E1 is ∼40 kb downstream of the *Peg13* DMR, and E2 is adjacent to the *Trappc9* intronic CTCF site that anchors the major maternal TAD (Fig. 6A). Both enhancers have high levels of H3K27ac and H3K4me1, and low levels of H3K4me3 in mouse brain (Fig. 6A). Moreover, an allelic H3K27ac dataset from hybrid mouse brain shows a maternal bias in enhancer activity. These regions do not show active enhancer marks in peripheral tissues (Fig. S4A), indicating they are brain-specific enhancers with preferential activity on the maternal allele. Additionally, both enhancers make maternally-biased contacts with *Kcnk9* in mouse brain (Fig. 6B), further supporting their potential role in driving maternal-specific expression of *Kcnk9* in neurons.

**Figure 6.**
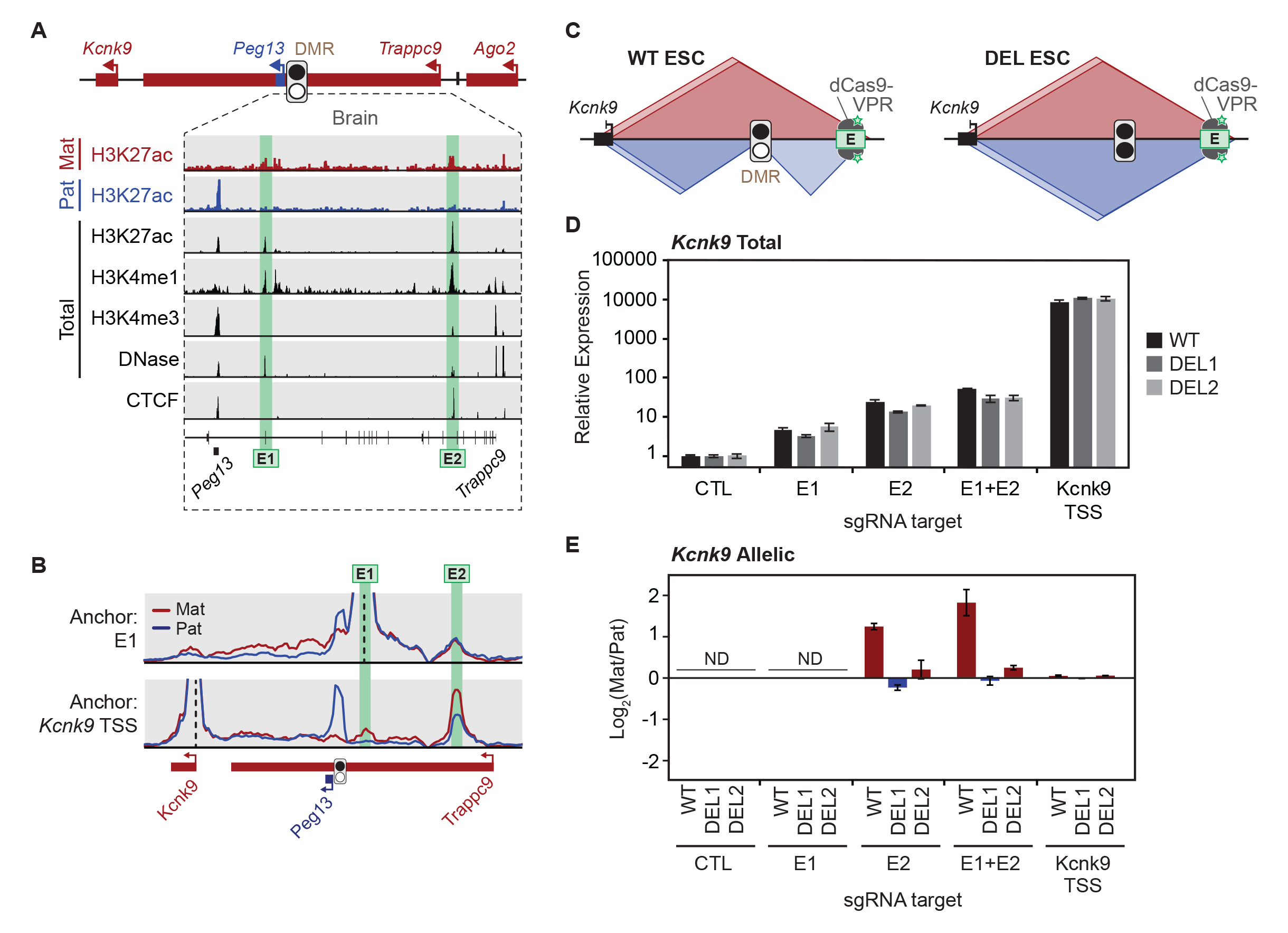
Pre-existing allelic chromatin structure primes imprinted expression upon enhancer activation. **A.** ChIP-seq genome browser tracks from mouse brain of histone modifications, DNA accessibility, and CTCF binding at E1 and E2. Total H3K27ac, H3K4me1, H3K4me3, and CTCF ChIP-seq are from ENCODE mouse postnatal day 0 whole brain samples and allelic H3K27ac is from Xie *et al*., 2012^4^. **B.** Virtual 4C from region capture Hi-C on hybrid mouse brain showing maternally biased contacts between the *Kcnk9* TSS and the enhancers. **C.** Diagram of CRISPR activation experimental design. dCas9-VPR was guided to the enhancers (E2 depicted) or the *Kcnk9* TSS. **D.** Total expression of *Kcnk9* as measured by RT-qPCR after guiding dCas9-VPR to the indicated target sites in WT and DEL ESCs. CTL samples were transfected with the sgRNA plasmid backbone with no sgRNAs inserted. **E.** Allelic expression of *Kcnk9* as measured by RT-ddPCR following dCas9-VPR treatment in WT and DEL ESCs. ND, not detected. See also Figure S4.

Given that allelic chromatin structure at the *Peg13*-*Kcnk9* locus is formed prior to the onset of *Kcnk9* expression, we hypothesized that pre-existing allelic contacts with these enhancers may be driving the maternal-specific imprinted expression of *Kcnk9* in neurons. We therefore asked if premature activation of the enhancers in the context of pre-existing allelic chromatin structure would drive maternal-specific expression of *Kcnk9*. We tested this in hybrid ESCs, which do not yet express *Kcnk9* but do have imprinted chromatin structure at the *Peg13- Kcnk9* locus. We guided biallelic transcriptional activation of the enhancers in WT ESCs using dCas9-VPR and multiplexed sgRNAs targeted to the enhancers (Fig. 6C). We then assessed the effects on gene expression in the *Peg13-Kcnk9* locus. We found that activation of E1 and E2 individually increased *Kcnk9* expression 5-fold and 25-fold, respectively. Activating both enhancers simultaneously increased *Kcnk9* expression over 50-fold (Fig. 6D). This upregulation occurred in a maternal-biased manner in WT ESCs (Fig. 6E). These results are not due to allelic differences in chromatin state at the *Kcnk9* promoter, as directly targeting the promoter of *Kcnk9* with dCas9-VPR led to a strong, biallelic upregulation of *Kcnk9* (Fig. 6D, E), and nanopore sequencing of the *Kcnk9* promoter showed that both alleles are unmethylated (Fig. S4B, C).

This supports a model in which both alleles of *Kcnk9* are accessible and transcriptionally competent, but only the maternal allele can be induced by enhancer activation due to differential chromatin structure. To further test this model, we then examined the effects of biallelic enhancer activation in DEL1 and DEL2 ESCs which do not have any imprinted chromatin structure. In this case, biallelic activation of the enhancers led to biallelic activation of *Kcnk9*, demonstrating that the allelic effect of enhancer activation is dependent on allelic chromatin structure (Fig. 6D, E). In contrast to the strong effects of enhancer activation on *Kcnk9* imprinted expression, the effect on the other genes in the locus was more modest, with only a slight upregulation of *Peg13* expression and no increase in *Trappc9* or *Ago2* expression (Fig. S4D-F), even though the promoters of these genes are hundreds of kilobases closer to the enhancer than the *Kcnk9* promoter. Overall, these results demonstrate that maternal-biased enhancer-promoter contacts can cause imprinted expression of *Kcnk9*, and that pre-existing allelic chromatin structure in ESCs primes the locus for imprinted expression upon enhancer activation.

## Discussion

In this study, we sought to characterize the cis-regulatory mechanisms underlying brain-specific imprinted expression of the *Peg13-Kcnk9* locus. We found that paternal-specific CTCF binding at the *Peg13* DMR leads to imprinted chromatin structure, and that this imprinted chromatin structure precedes imprinted gene expression at the locus. Additionally, we found that enhancer activation is sufficient to drive maternal-specific expression of *Kcnk9* in a manner that is dependent on pre-existing allelic chromatin structure.

We also tested a model in which the *Peg13* lncRNA itself acts *in cis* to repress the surrounding genes on the paternal allele. Other imprinted lncRNAs, such as *Kcnq1ot1*, *Airn*, and *Ube3a-ATS,* mediate gene repression through mechanisms involving recruitment of cis-repressive factors^15,16,51,52^ and transcriptional interference.^17,57,58^ We found that knockdown of the *Peg13* lncRNA does not affect the expression of surrounding genes, suggesting *Peg13* is not a cis-repressive lncRNA. Other recent work has attempted to elucidate the function of *Peg13* and has proposed that it may act as a sponge for various microRNAs^56,59,60^ and suppress *Yy1* through the recruitment of the PRC2 complex.^53^

In addition to its role in mediating allelic chromatin structure, our data also suggest that CTCF is important for maintaining an open chromatin state at the paternal *Peg13* DMR. Deleting the CTCF binding sites at the DMR led to a gain of methylation of the surrounding region, which is similar to previous observations at the *H19-Igf2* locus.^34^ ICR maintenance is an area of ongoing research and is not fully understood. DNA-binding proteins such as ZFP57 and TRIM28 are important for protecting ICRs from the wave of global demethylation that occurs during early embryogenesis.^13,14,61,62^ CTCF is bound to the unmethylated allele of many ICRs and DMRs,^46^ and it may play a part in ICR maintenance across multiple imprinted regions in addition to its role in higher-order chromatin organization.

While this work provides important insights into brain-specific imprinted expression of the *Peg13-Kcnk9* locus, several key questions remain. It remains to be determined how the imprinted biases of *Trappc9*, *Chrac1*, and *Ago2* are acquired during neuron differentiation. We did not observe a change in *Trappc9* or *Ago2* levels upon enhancer activation or *Kcnk9* TSS activation, suggesting that the mechanism underlying *Trappc9* and *Ago2* imprinted expression may be different from that of *Kcnk9*. While the imprinted expression of *Kcnk9* and *Peg13* seen in mouse is known to be conserved in human,^39^ the maternal bias of *Trappc9* and *Ago2* may not be present in human brain,^39^ further supporting the notion that *Trappc9* and *Ago2* imprinting could be achieved through a different mechanism than *Kcnk9*.

Genetic mutations in the *Peg13-Kcnk9* locus are associated with intellectual disability,^40,41,42,43^ and our current work on the mechanistic basis of imprinting at the *Peg13- Kcnk9* locus may be informative for future therapeutic approaches. Previously, re-activation of the normally silent paternal allele of *Kcnk9* by global histone deacetylase inhibition rescued the behavioral phenotype in a Birk-Barel syndrome mouse model, a disease caused by a missense mutation in maternal *Kcnk9*.^63^ In our study, we found that the paternal allele of *Kcnk9* was reactivated upon deletion of the CTCF-containing region of the DMR, highlighting additional therapeutic opportunities in Birk-Barel syndrome, for example through dCas9-mediated blocking of CTCF at the paternal DMR.

The role that higher-order chromatin structure plays in gene regulation remains a matter of debate. Many lasting features of chromatin structure are established genome-wide early in development.^2^ But dynamic changes in chromatin contacts during development correlate with transcriptional changes,^64^ leading to questions about the cause-effect relationship of chromatin structure and gene expression. Deletion of individual CTCF sites often leads to changes in expression levels of nearby genes,^25,26,27^ but global depletion of CTCF causes differential expression of a relatively small number of genes.^28,29^ Genomic imprinting has provided a crucial model system for the exploration of these questions, and will likely continue to do so in the future.

Our data suggests a model whereby chromatin structure established early in development can prime lineage-specific gene expression patterns. Whether this mechanism will apply to other loci with tissue-specific imprinted expression remains to be seen. Our finding that allelic chromatin structure at the *Peg13-Kcnk9* locus is largely static during differentiation is consistent with recent experiments on the *Dlk1-Dio3* imprinted locus, where the paternal-specific activation of *Dlk1* upon neurogenesis occurs in the absence of corresponding changes in chromatin structure on the paternal allele.^20^ It will be interesting to see if this holds true for genes with more complex tissue-specific imprinted expression patterns. *Igf2* is known to be paternally expressed in early development in a chromatin-structure dependent manner, but it intriguingly switches to a maternal bias in the brain.^8^ Whether this happens in a static or dynamic chromatin structure environment, or whether this switch is dependent on chromatin structure at all, remains an open question. Another example is found at the *Grb10-Ddc* locus.

*Grb10* is maternally expressed in most adult tissues in mouse, but in neuronal tissue it is expressed from an alternative paternal-specific promoter.^65,66^ The maternal-specific expression in heart and muscle is depends on allelic chromatin structure due to allelic CTCF binding at a secondary DMR,^21^ but the role of chromatin structure in the switch to paternal expression in neuronal lineages has not been investigated. Overall, our work contributes to an emerging model that imprinted chromatin structure may be a widespread mechanism through which epigenetic inheritance regulates gene expression across tissues.

## Acknowledgements

We thank the Bauer Core Facility at Harvard University for technical support and access to instrumentation, including high-throughput sequencing, flow cytometry, and ddPCR. This work was supported by the Klingenstein-Simons Fellowship Award in Neuroscience (A.J.W.), the George W. Merck Fellowship Fund (A.J.W.), the Rita Allen Foundation (A.J.W.), and National Institute of General Medical Sciences of the National Institutes of Health under Award Number R35GM146921 (A.J.W.). The content is solely the responsibility of the authors and does not necessarily represent the official views of the National Institutes of Health.

## Author Contributions

Conceptualization and methodology, D.L. and A.J.W.; Investigation, D.L., C.M.W., B.B.; Formal Analysis, D.L.; Writing-Original draft: D.L.; Writing-Review and editing, A.J.W.; Supervision and funding acquisition: A.J.W.

## Supplementary Figure Legends

**Figure S1.**
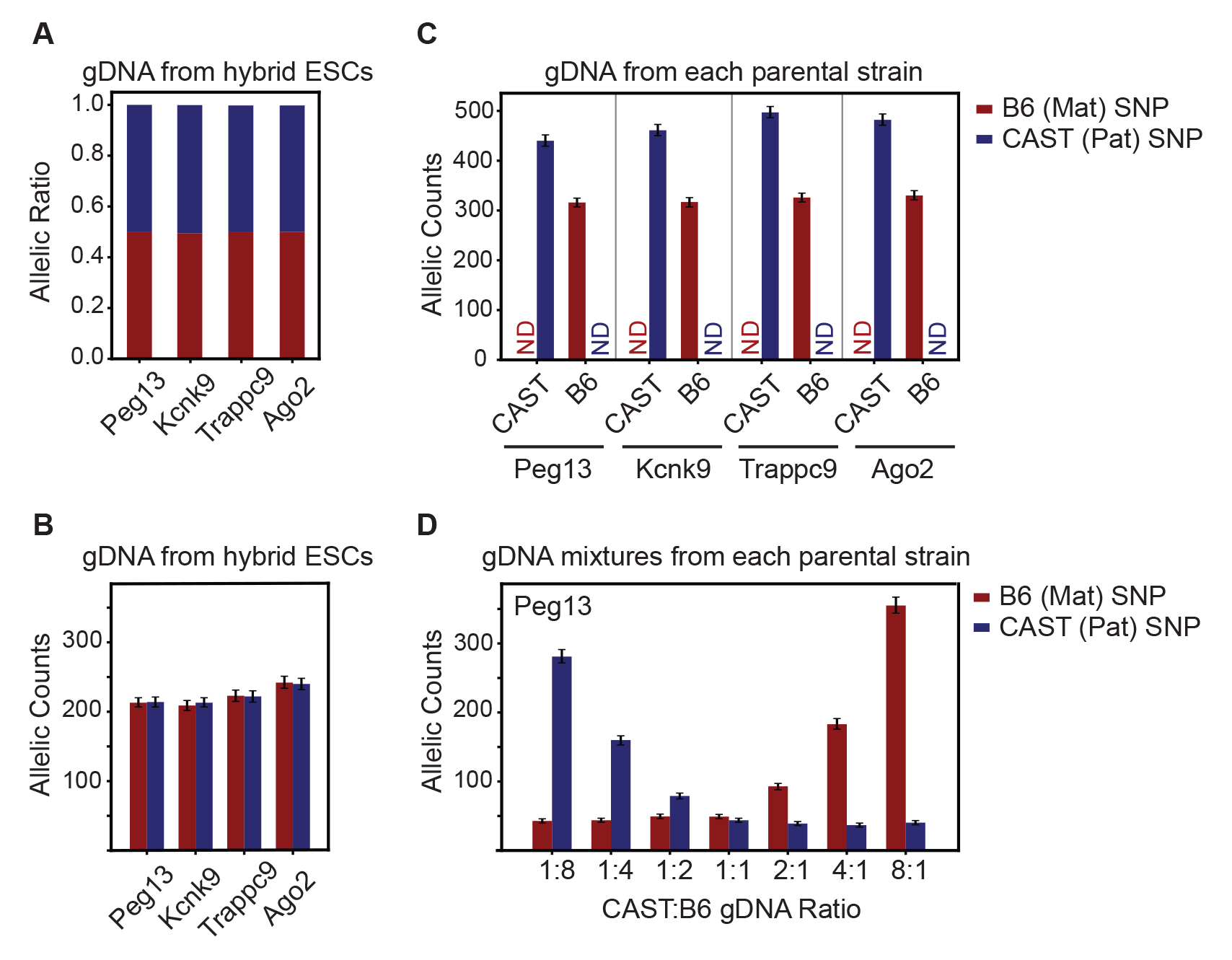
Confirmation of allelic quantification by ddPCR, related to Figure 1. **A-B.** Allelic ddPCR for each of the indicated genes on genomic DNA from hybrid ESCs showing maternal to paternal allelic ratio (A) and allelic counts (B). **C.** Allelic ddPCR for each of the indicated genes on genomic DNA from B6 or CAST strains. ND, not detected. **D.** Allelic ddPCR for *Peg13* on genomic DNA from B6 or CAST strains mixed at the indicated ratios. Error bars, Poisson 95% confidence interval.

**Figure S2.**
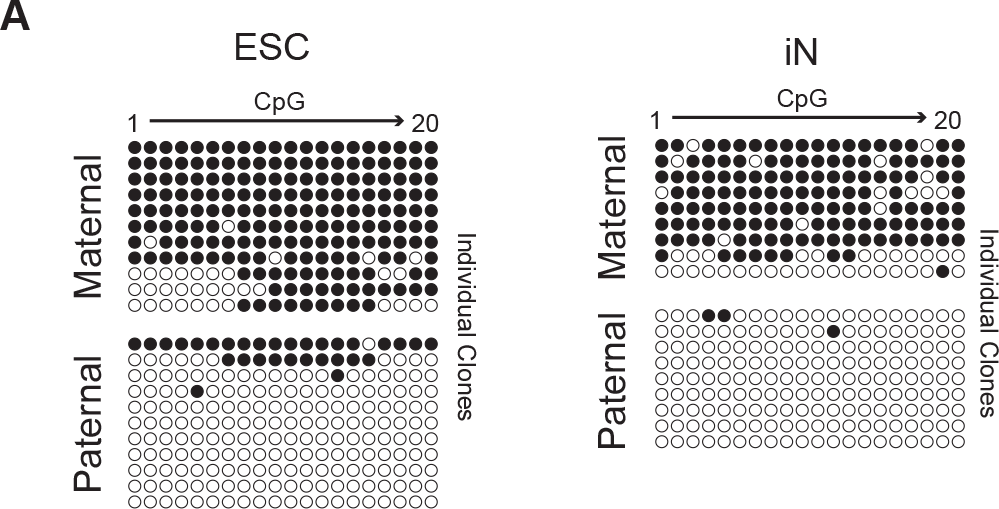
Bisulfite Sanger-sequencing of the *Peg13* DMR, related to Figure 2. **A.** Bisulfite Sanger-sequencing in ESCs and iNs indicating methylated (closed circles) and unmethylated (open circles) CpGs.

**Figure S3.**
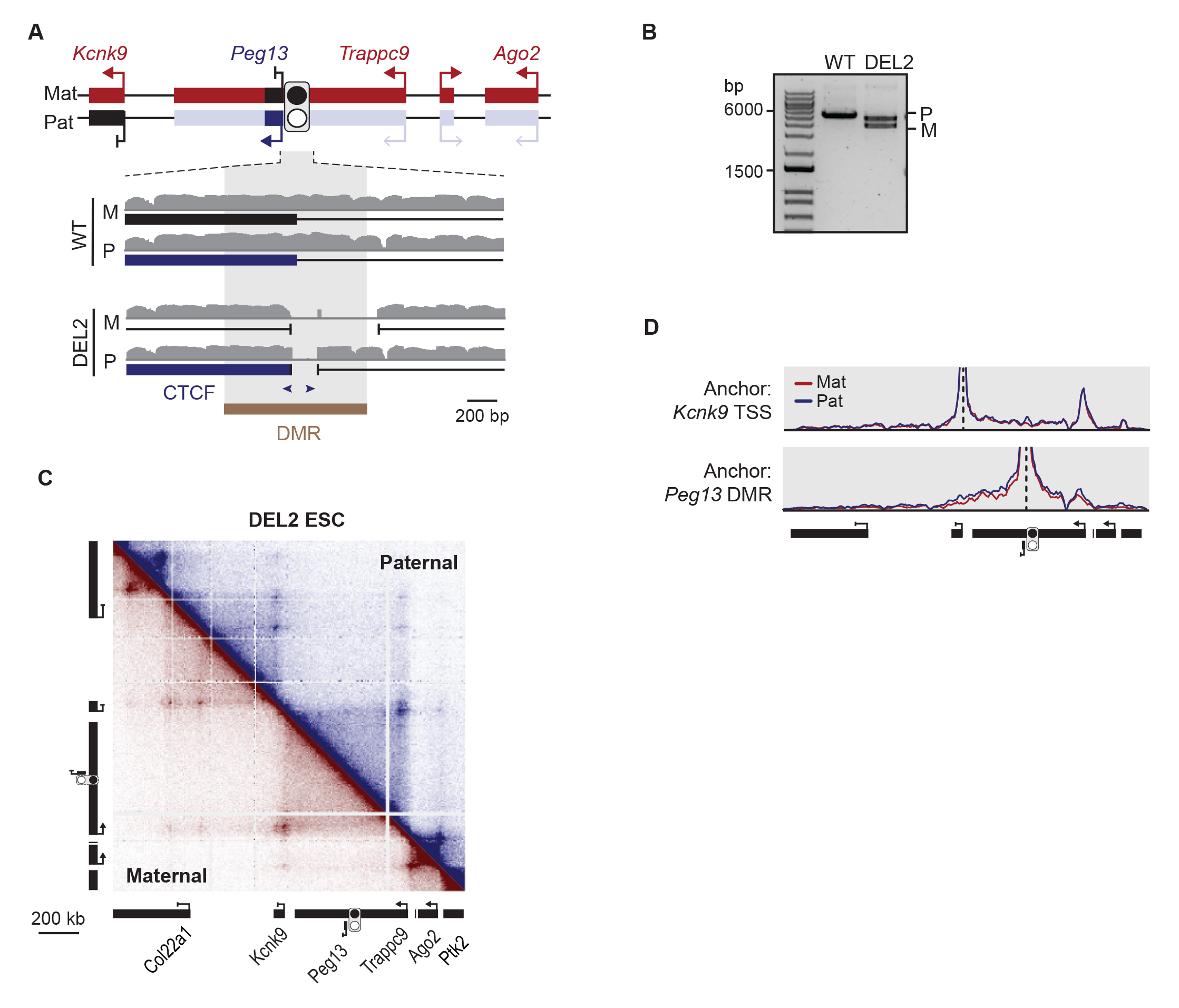
Characterization of DEL2 chromatin structure, related to Figure 4. **A.** Diagram of the *Peg13* DMR depicting the CRISPR-mediated deletions of DEL2. Genomic tracks show single-end mapping of Hi-C genomic DNA. **B.** Agarose gel showing PCR products from the clonal ESC line using primers spanning the *Peg13* DMR. Deletion products were validated by Sanger sequencing to determine parent-of-origin. **C.** Allelic region capture Hi-C of DEL2 ESCs. **D.** Virtual 4C analysis of DEL2 ESC region capture Hi-C.

**Figure S4.**
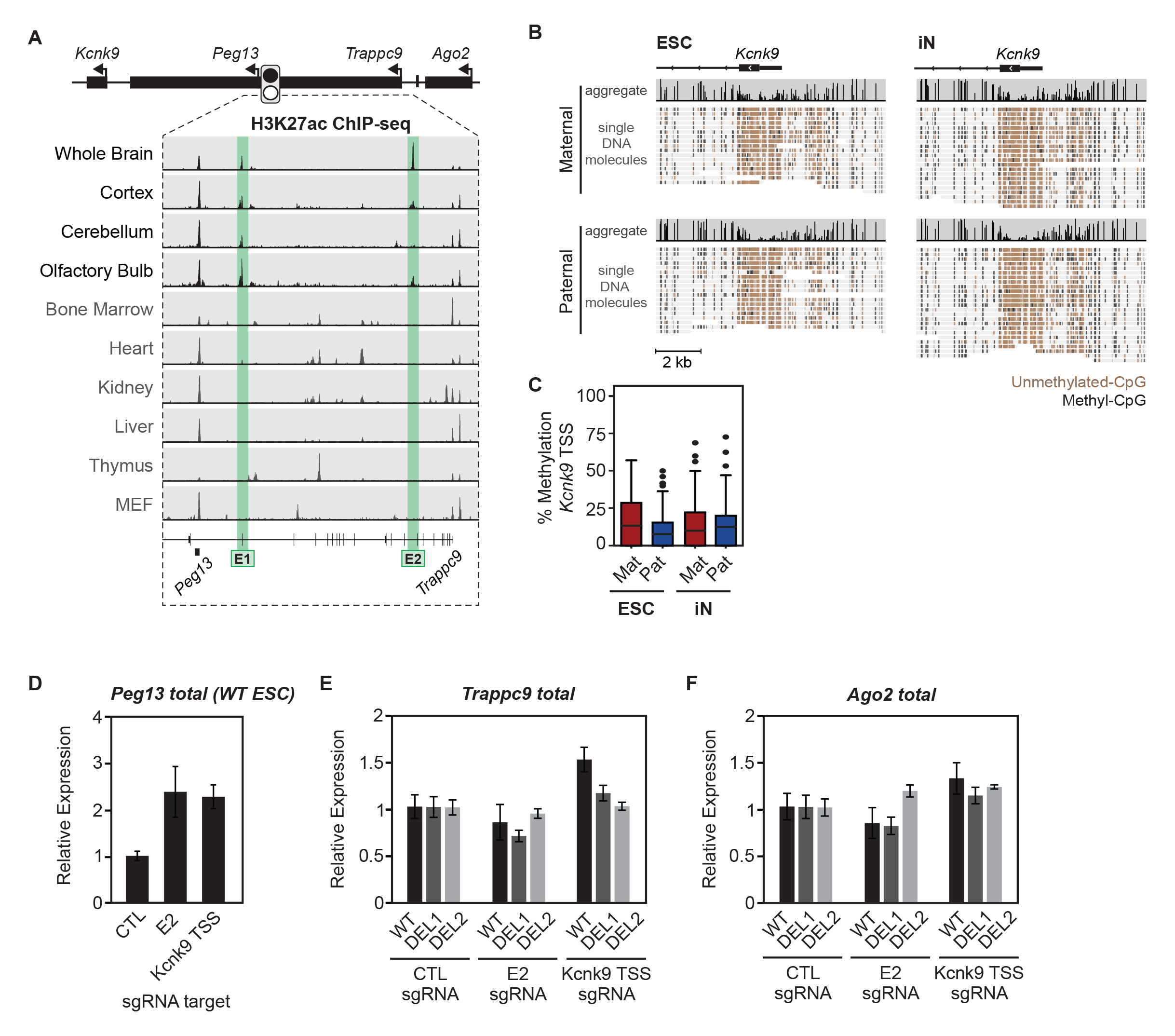
Enhancer activation gene expression profiles, related to Figure 6. **A.** ENCODE tracks across mouse tissues for H3K27ac. **B.** Targeted nanopore sequencing of native genomic DNA of the *Kcnk9* TSS to determine CpG methylation levels in WT ESCs and iNs. **C.** Quantification of (B). **D.** Total *Peg13* expression as measured by RT-qPCR after guiding dCas9-VPR to the indicated target sites in WT ESCs. Not that this was not done in DEL ESCs as the *Peg13* promoter is deleted in those cell lines. CTL samples were transfected with the sgRNA plasmid backbone with no sgRNAs inserted. **E-F.** Total expression of *Trappc9* (E) *and Ago2* (F) as measured by RT-qPCR after guiding dCas9-VPR to the indicated target sites in WT and DEL ESCs.

**Table S1.**
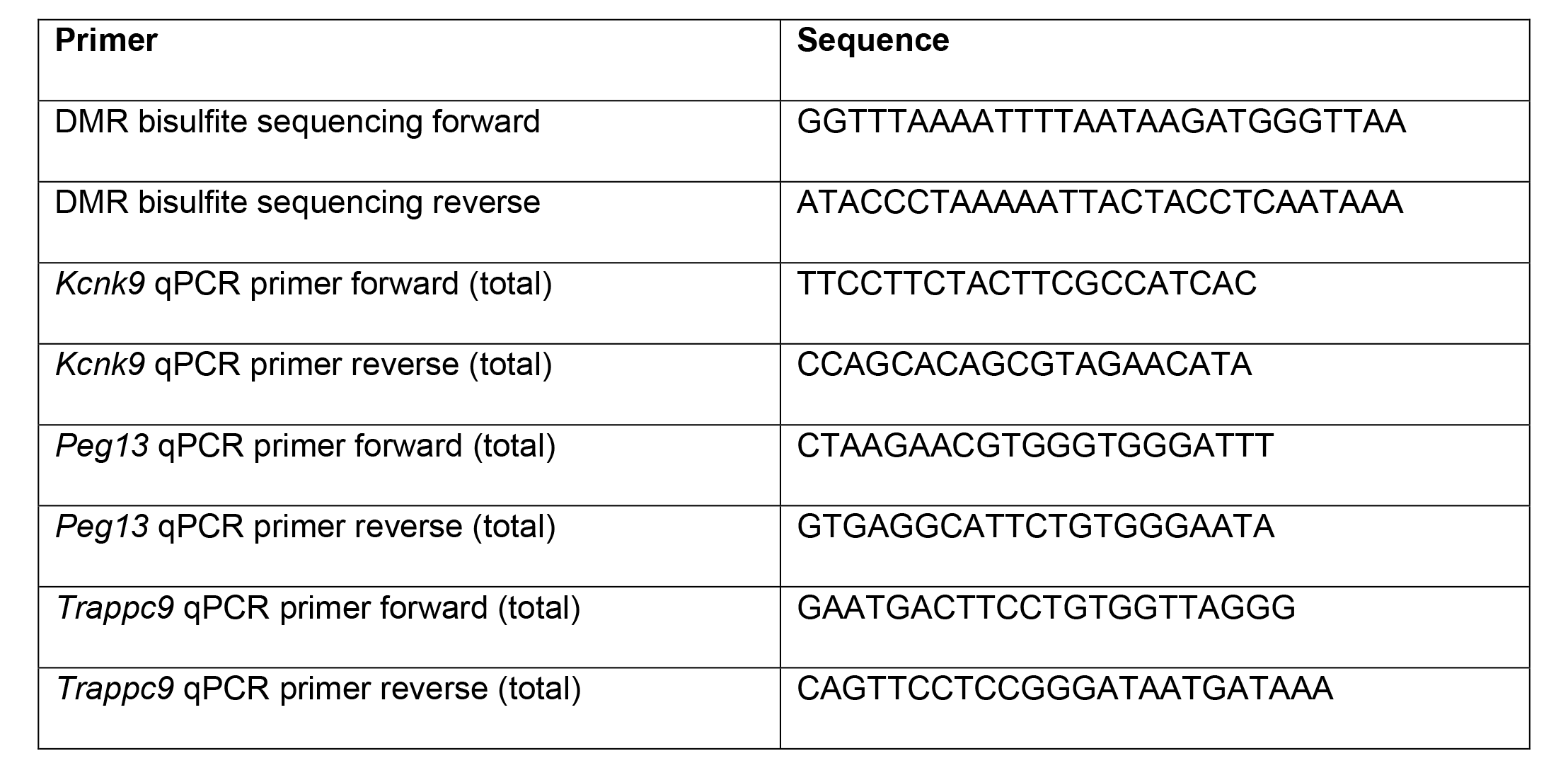
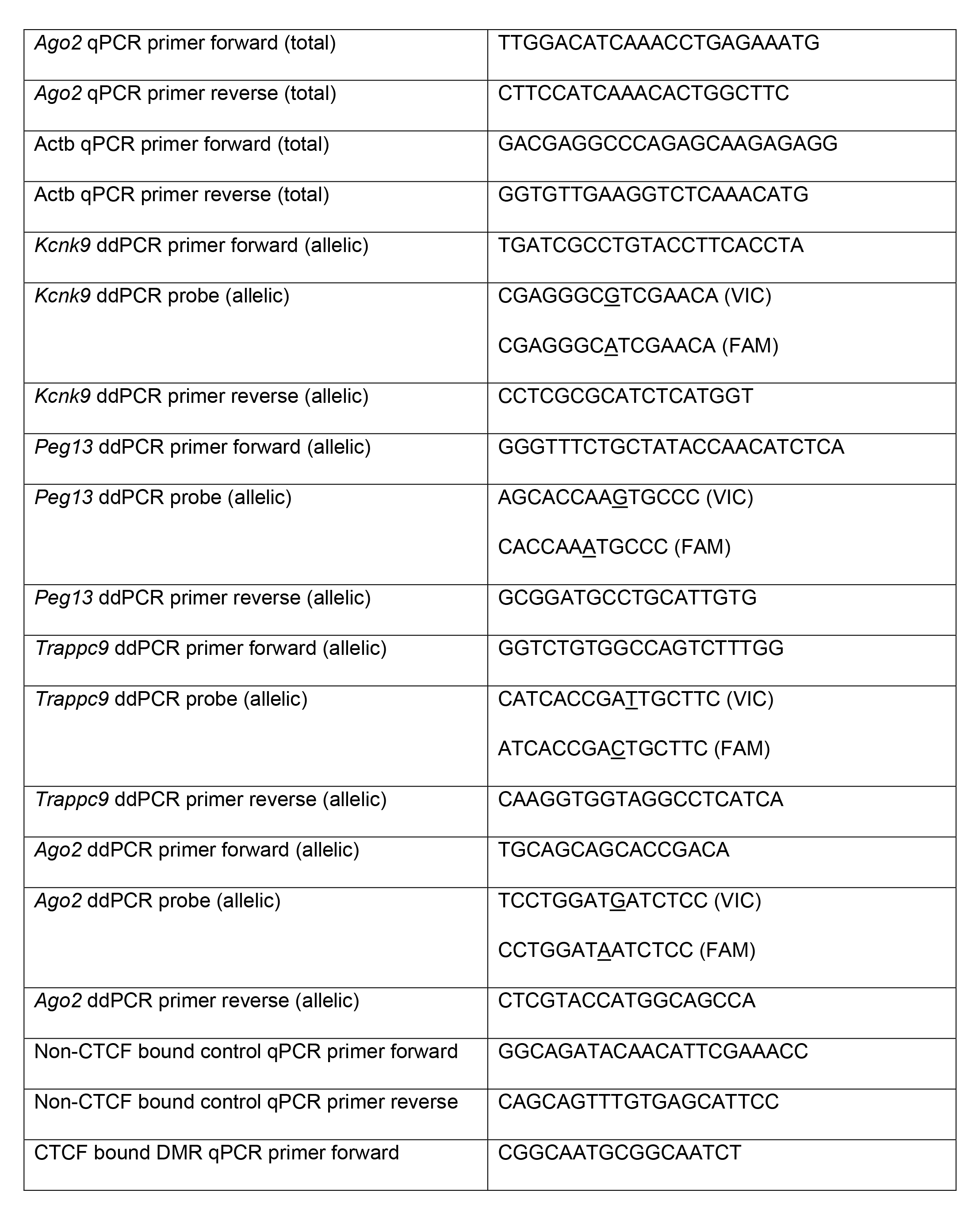
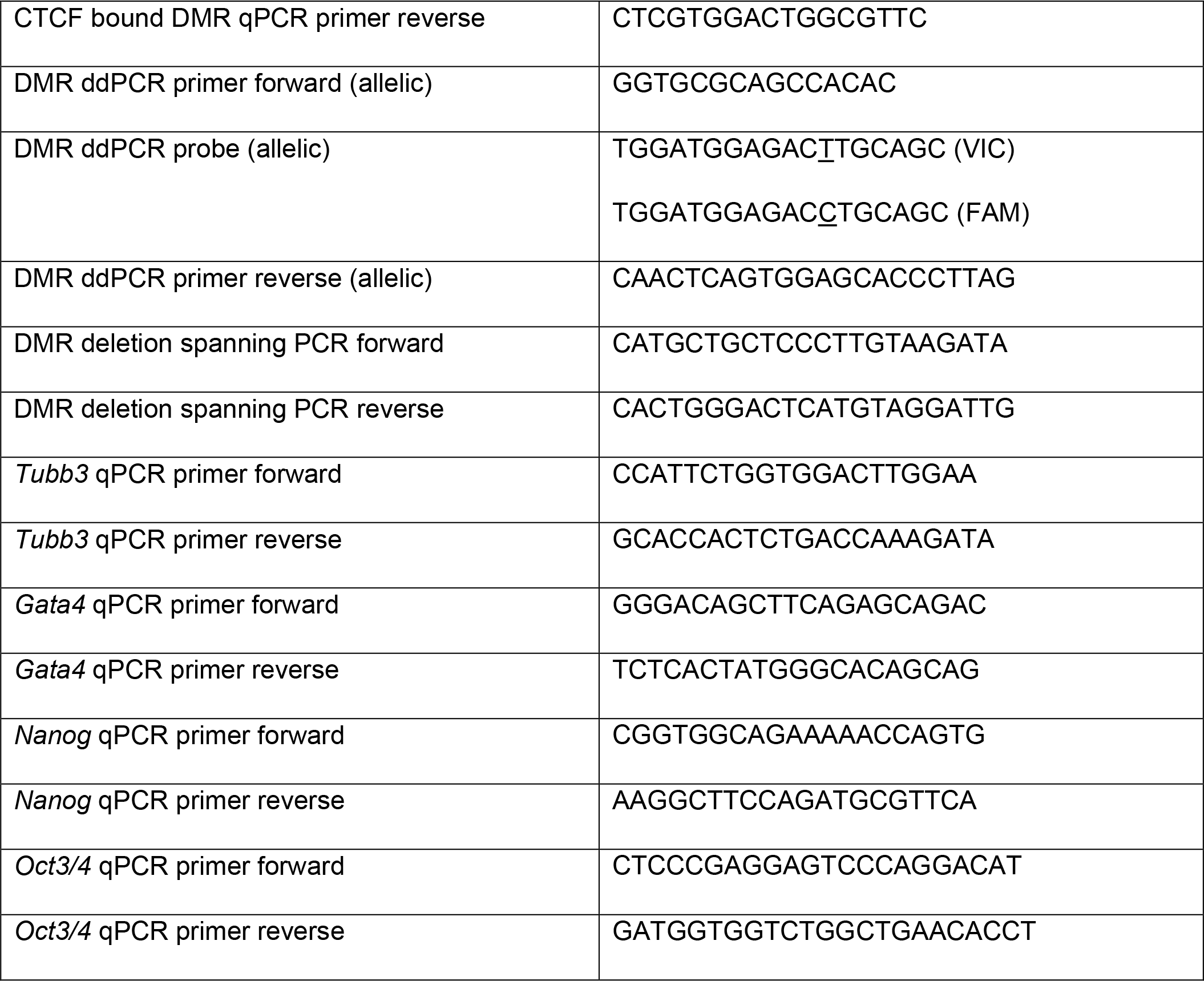
PCR primer sequences, related to Methods.

**Table S2.**
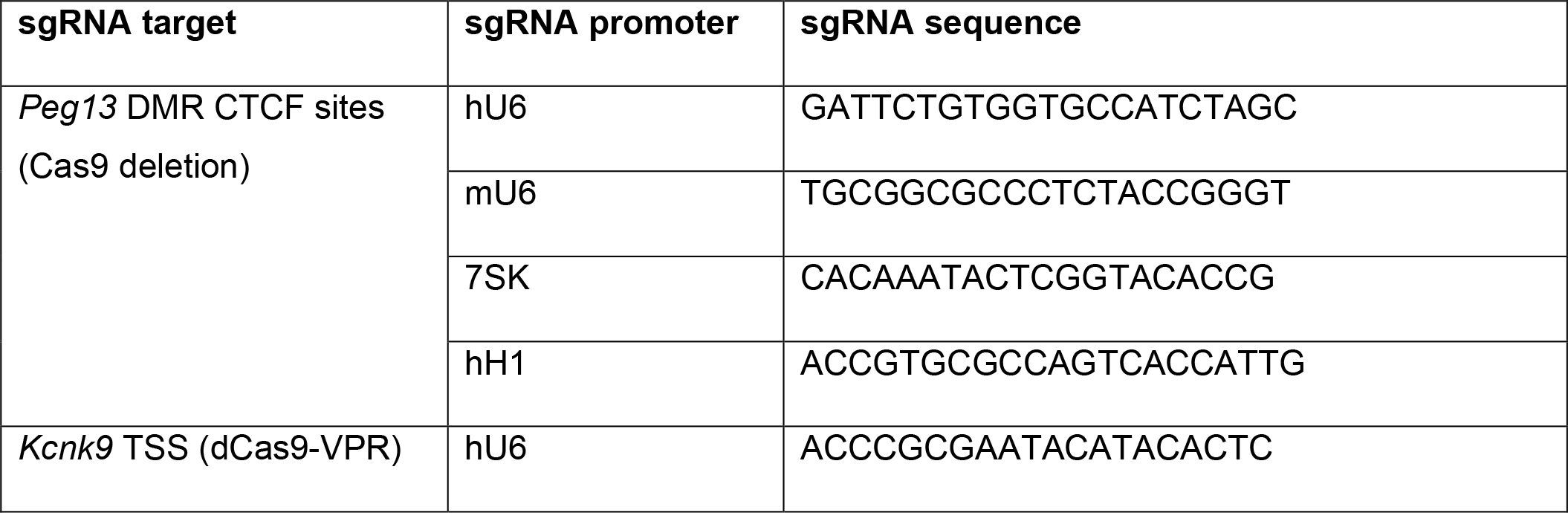
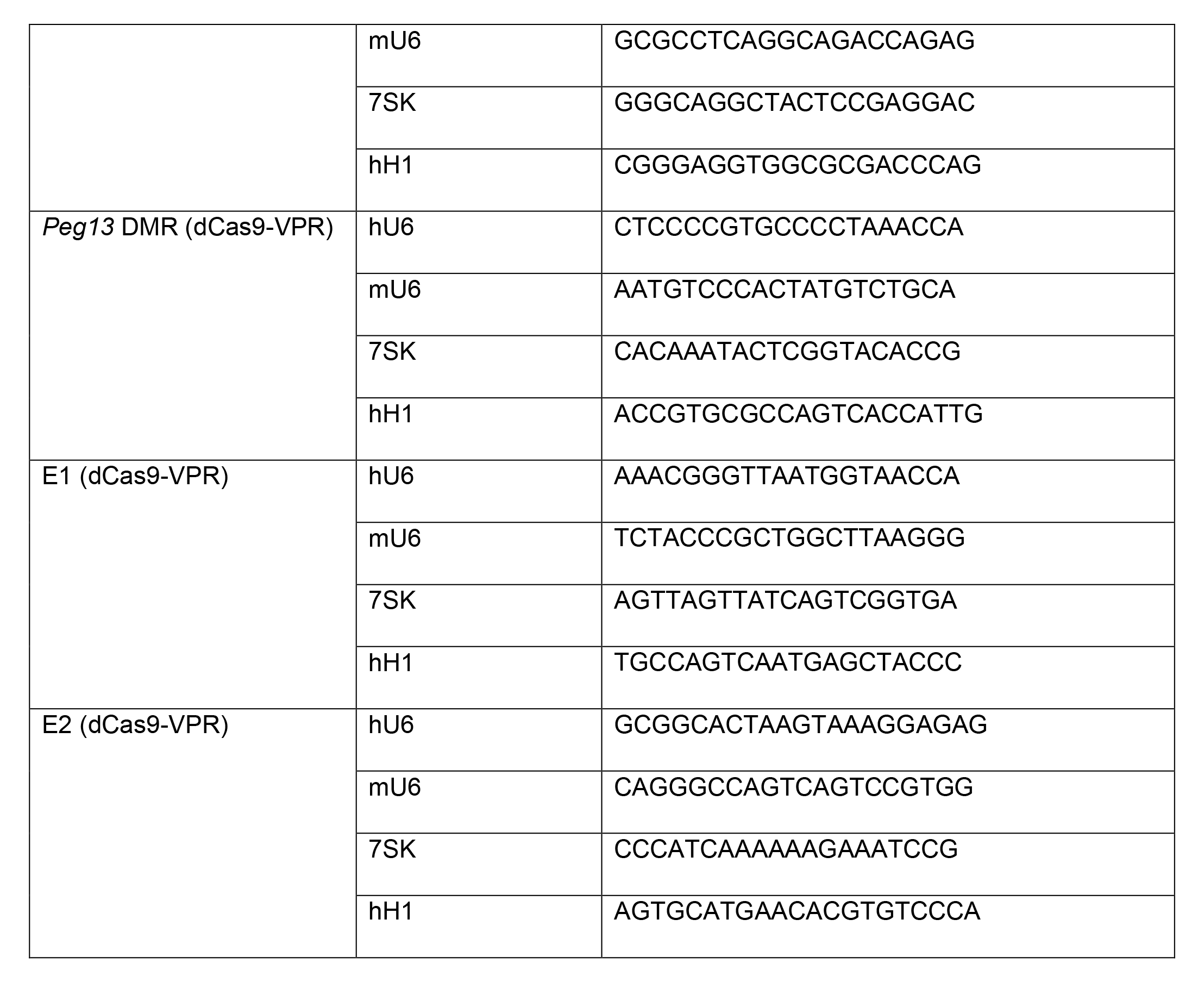
sgRNA sequences, related to Figure 6.

## Methods

### ESC line generation and culture

The ESC line used in this study were derived from an F1 *Mus. Musculus* (129/Sv) x *Mus. Castaneous* cross, F1.2-3.^67^ Two PiggyBac plasmids were co-transfected into ESCs for stable integration of EF1α-rtTA (PB-EF1α-M2rtTA;CMV-HygroR) and TetO-Ngn2 (PB-TetO-Ngn2;SV40-NeoR) in the presence of piggyBac transposase. ESCs with the integrated constructs were selected for 10 days in ESC media containing Hygromycin B (150 µg/ml) and Geneticin (300 µg/ml). A single cell ESC clonal line (F1.2-3; EF1α-rtTA;TetO-Ngn2 clone D4) with high differentiation potential and maintenance of proper imprinting at the *Peg13*-*Kcnk9* locus was selected for all studies described here. To passage ESCs, cells were washed with HEPES-buffered saline (HBS), dissociated using 0.25% Trypsin in HBS with 1 mM EDTA, and then transferred onto plates pre-coated with 0.2% gelatin in phosphate-buffered saline (PBS). ESCs were maintained in serum/LIF ESC media composed of Dulbecco’s Modified Essential Media (Thermo) supplemented with 10 mM HEPES (Thermo), 0.11 mM β-mercaptoethanol (Sigma), 1X nonessential amino acids (Corning), 2 mM L-glutamine (Corning), 1X penicillin streptomycin solution (Corning), 15% fetal bovine serum (Cytivia), and 1000 U/mL leukemia inhibitory factory (Sigma). Cells were maintained in a humidified 5% CO_2_ incubator at 37°C and split every 48 hours or when cells reached 70% confluency, whichever came first.

### Neuron differentiation

ESCs were cultured for 24 hours on gelatin-coated plates in ESC media (as above), then ESCs were washed and cultured for 24 hours in neuron differentiation media [BrainPhys media supplemented with 1X SM1 neuronal supplement (STEMCELL Technologies), 1X N-2 supplement (Thermo), 20 ng/mL brain-derived neurotrophic factor (STEMCELL Technologies), and 1 µg/ml doxycycline (Sigma)]. Day 1 iNs were then dissociated using accutase (STEMCELL Technologies), counted, and plated on PEI-coated plates (0.1% PEI in borate buffer containing 50 mM boric acid and 24 mM sodium tetraborate at pH 8.4) in neuron differentiation media at a density of 40,000 cells/cm^2^. Half of the media was replaced every 48 to 72 hours for a total of six days, unless otherwise noted.

### RNA isolation and cDNA synthesis

Total RNA was harvested from ESCs or iNs in biological triplicate using TRIzol (Thermo) and subject to DNase treatment using TURBO DNase (Thermo) (Fig. 1 and 4) or using the Zymo Quick-RNA Microprep Kit with on-column DNase treatment (Fig. 3). Reverse transcription was performed using SuperScript III Reverse Transcriptase (Thermo) with random hexamers.

### qPCR

qPCR reactions were prepared using Power SYBR Green (Thermo) and run on the Bio-Rad CFX Opus 384 Real-Time PCR System. Relative expression was calculated using the delta-delta Ct method relative to ActB.

### ddPCR

PCR primer/probe sets were designed using the Thermo Custom Taqman SNP Genotyping Assay Design Tool to overlap a *musculus*/*castaneous* SNP. Probes overlapping the *musculus* (maternal) or *castaneous* (paternal) SNP were VIC-conjugated or FAM-conjugated, respectively. Samples were prepared using ddPCR Supermix for Probes (No dUTP) (Bio-Rad). Droplets for ddPCR were generated using the Bio-Rad QX200 Droplet Generator, PCR was performed according to the Supermix protocol, and then droplets were read on the Bio-Rad QX200 Droplet Reader.

### Bisulfite sequencing

Genomic DNA was extracted from cell pellets using the Monarch gDNA Purification Kit (NEB). 2 µg of purified genomic DNA was then bisulfite converted using the EpiTect Bisulfite Kit (Qiagen). PCR was performed on bisulfite-treated gDNA using HotStarTaq DNA polymerase (Qiagen). PCR product was then TOPO cloned (Thermo) and transformed into NEB Stable Competent *E. coli*. Individual colonies were picked and subject to Sanger sequencing of the *Peg13* DMR (see Table S1 for primer sequences). SNPs were used to identify maternal and paternal DNA fragments, and sequencing traces were analyzed using BISMA^68^ to determine the methylation status of individual CpGs.

### Long-read methylome analysis

Isolated nuclei were subjected to GpC methylation using M.CviPI GpC methyltransferase (NEB), followed by proteinase K and RNaseA treatment. Genomic DNA was purified from the reaction using AMPure XP beads. A total of 6-12 µg of genomic DNA was dephosphorylated with rSAP (NEB), followed by AMPure XP beads purification. *In vitro* Cas9 incubation was performed to enrich the nanopore library for regions of interest using EnGen sgRNA synthesis kit (NEB) and Cas9 nuclease (NEB). Cleaved genomic DNA product was treated with NEBNext dA-tailing kit (NEB) and subjected to library preparation using nanopore adapter (ONT SQK-LSK109) and Quick T4 DNA ligase (NEB). Libraries were sequenced in MinION flow cell (ONT R9.4.1). Sequencing data was then mapped to the *castaneous* SNPs incorporated mouse genome using minimap2 (2.24).^69^ Variant tagging and haploid phasing of each sequencing read were performed using nanopolish (0.13.2)^70^ and whatshap.^71^ Mapped data was split into each haplotype using bamtools,^72^ and CpG methylation analysis was performed using nanopolish call-methylation package. Data was visualized in IGV.^73^

### Chromatin immunoprecipitation (ChIP)

Chromatin for ChIP was prepared and sheared using the truChIP Chromatin Shearing Kit (Covaris). Cells (∼5 million ESCs or iNs per biological replicate) in culture plates were fixed in 1% methanol-free formaldehyde (Thermo) at room temperature for 10 minutes and then quenched with glycine. Chromatin prepared using the truChIP kit was sheared to 500 bp in a volume of 130 µl using the Covaris S220 sonicator. Immunoprecipitation of chromatin was performed as follows. For each IP, 50 µl of protein A beads (Thermo) were prepared by washing on a magnetic stand three times using 0.1 M Na-phosphate buffer, pH 8. Beads were then suspended in 50 µl Na-Phosphate buffer and incubated at room temperature for 10 minutes with desired antibody [20 µl of CTCF antibody (Cell Signaling Technology #2899) or 11 µg of anti-rabbit IgG (Thermo)]. Antibody-conjugated beads were washed three times with Na-phosphate buffer and resuspended in 50 µl of Na-phosphate buffer. Resuspended beads were then added to 100 µl of sheared chromatin and rotated at 4°C overnight. Beads were then washed at 4°C, as follows: twice using low salt buffer (20 mM HEPES pH 8.0, 150 mM NaCl, 0.1% Triton X-100, 0.1% SDS, 2 mM EDTA), once using high salt buffer (20 mM HEPES pH 8.0, 500 mM NaCl, 0.1% Triton X-100, 0.1% SDS, 2 mM EDTA), twice using LiCl buffer (100 mM Tris-HCl pH 7.5, 0.5 M LiCl, 1% NP-40, 1% Na-deoxycholate), and twice using TE buffer (10 mM Tris pH 8.0, 1 mM EDTA). Beads were then resuspended in 250 µl elution buffer (10 mM Tris pH 8.0, 1 mM EDTA, 1% SDS) and incubated at 65°C for 30 minutes to elute. Samples were then pelleted and the supernatant was transferred to a new tube and incubated overnight at 65°C to reverse crosslinking. 250 µl of TE buffer was added to each IP, and then samples were treated with 0.2 µg/µl RNase A at 37°C for 1 hour followed by 0.2 µg/µl proteinase at 55°C for 1 hour. DNA was then purified using phenol-chloroform extraction followed by ethanol precipitation and subject to qPCR or ddPCR analysis.

### Region capture Hi-C

Hi-C libraries were generated using the Arima-HiC Kit with 3 µg of chromatin from ESCs, iNs, or cortical quarters as input. End repair and adapter ligation were performed using the Accel-NGS 2S DNA Library Kit (Swift Biosciences). Agilent SureSelect probes against the *Peg13-Kcnk9* locus were designed against mm10 chr15:71850001-73350000. Probes overlapping *musculus*/*castaneous* SNPs were designed to perfectly match both genotypes (i.e. two probes per SNP) for *in vitro* experiments and B6 alone for *in vivo* experiments. End repaired and adapter ligated Hi-C libraries were then enriched using Agilent SureSelectXT HS Target Enrichment System. Enriched DNA was amplified and sequenced on Illumina NextSeq Sequencing System.

### Region capture Hi-C analysis

The mm10 reference genome was N-masked using SNPsplit_genome_preparation from SNPsplit.^74^ The N-masked reference genome was then digested *in silico* according to the Arima restriction enzymes using HiCUP.^75^ Bowtie2^76^ was then used to map fastq files to the digested N-masked reference genome using the HiCUP wrapper. Resulting SAM files were split into maternal and paternal reads using SNPsplit. Maternal and paternal SAM files from biological replicates and reciprocal crosses were merged by parent of origin by randomly subsampling higher read depth samples such that an equal number of reads from all replicates and crosses were represented in the final merged file. Allelic SAM files were converted to the Juicer medium format^77^ and then converted to .hic format using Juicer pre. Final Hi-C plots were visualized using Juicebox at 5 kb resolution.^78^ Allelic subtraction plots were generated using Juicebox. Virtual 4C plots were generated by summing the contacts made from a 10 kb anchor point overlapping the genomic feature of interest and all surrounding genomic regions using a 5 kb sliding window at 1 kb resolution. Insulation score was determined by summing all contacts made in a sliding 200 kb window at 10 kb resolution.

### Generation of sgRNA vectors

dCas9-VPR and Cas9 experiments were performed using a published multiplex four sgRNA construct system.^79^ The four sgRNAs for each of the sgRNA target sites (*Kcnk9* TSS, E1, and E2) were cloned into four sgRNA backbones each containing different small RNA promoters (Addgene #53186, #53187, #53188, #53189). Golden Gate cloning into pLV-GG-hUbC-dsRED (Addgene #84034) generated the final vector co-expressing four sgRNAs.

### CRISPR deletion of DMR CTCF region

ESCs were co-transfected with 1000 ng of pX458 SpCas9(BB)-2A-GFP vector (Addgene #48138, contains eGFP)^80^ and 500 ng of vector co-expressing four sgRNAs to the *Peg13* DMR CTCF region (contains DsRed) mixed with 3.75 µl of Lipofectamine 2000 (Thermo) in 500 µl of Opti-MEM (Thermo). The mixture was then plated into a suspension of 500,000 ESCs in 1.5 ml ESC media into individual wells of a gelatin-coated six-well culture plate. After 24 hours, fresh ESC media was added. The next day, eGFP+/DsRed+ cells were bulk sorted by FACS and expanded. Subsequently, single cells were sorted to establish clonal lines and expanded. Genomic DNA was extracted using QuickExtract (Epicentre) and PCR amplified across sgRNA sites (Table S1). PCR bands corresponding to deletion products were gel isolated and subject to Sanger sequencing. SNPs in the amplicon allowed for allelic determination of the deletion products. Wild-type clonal ESC lines were generated from the same procedure performed in the absence of sgRNA (i.e. no sgRNA vector).

### CRISPR activation with dCas9-VPR

2 µg total of multiplex sgRNA vector and EF1α-dCas9-VPR-Puro vector (Addgene #99373) at a 2:1 molar ratio were mixed with 5 µl of Lipofectamine 2000 (Thermo) in 500 ul of Opti-MEM (Thermo). The mixture was then plated into a suspension of 600,000 ESCs in 1.5 ml ESC media into individual wells of a gelatin-coated six-well culture plate. Media was then changed after 24 hours to fresh ESC media. The next day, media was changed to ESC media containing 1 µg/ml puromycin. After 24 hours of drug selection, ESCs were washed and RNA was purified using TRIzol as above and subject to qPCR or ddPCR analysis. As a negative control, ESCs were generated from the same procedure performed in the absence of sgRNA (i.e. blank sgRNA backbone vector).

### Analysis of allelic ChIP-seq data

To analyze previously published allelic ChIP-seq datasets from hybrid mouse brain, raw fastq files were downloaded from GSE35140^46^ (CTCF) or GSE33722^4^ (H3K27ac). Reads were trimmed using TrimGalore^81^ and then aligned using STAR^82^ to mm10 that had been N-masked using SNPsplit. Reads were then split into maternal and paternal reads using SNPsplit as above. Resulting BAM files were then converted to bigwigs using deepTools bamCoverage^83^ and visualized in IGV.

### Single molecule FISH

RNAScope was performed on 16 µm frozen adult mouse brain sections. Sections were fixed in cold 4% paraformaldehyde for 15 minutes. Sections were then progressively dehydrated in 50% ethanol, 70% ethanol, and 100% ethanol for 5 minutes each. Single molecule FISH was then performed using the RNAScope Fluorescent Multiplex Kit (ACD Bio). Sections were counterstained with DAPI and cover slipped with Aqua-Poly/Mount (Polysciences). Tissue sections were imaged on an Axioscan 7 (Zeiss) microscope with a 64X objective.

## Data and code availability

This paper does not report original code. Raw and processed region capture Hi-C data have been deposited on GEO under accession number GSE202251.

